# ChatDIA: A zero-shot large language model workflow for targeted analysis of data-independent acquisition mass spectrometry data

**DOI:** 10.64898/2026.02.11.705360

**Authors:** Jiayi Li, Joshua Charkow, Mingxuan Gao, Jiaqing Li, Hannes Röst

## Abstract

Data-independent acquisition (DIA) proteomics enables reproducible, large-scale protein identification and quantification but remains challenging to analyze due to highly complex MS/MS spectra and chromatographic interference, particularly in low signal-to-noise single-cell proteomics. Here, we introduce ChatDIA, a zero-shot large language model (LLM)-based workflow for targeted DIA analysis that operates through an explicit reasoning-based decision framework. ChatDIA performs automated peptide identification and supports natural-language interaction with DIA data. Unlike purpose-built DIA software that relies on domain-specific models, ChatDIA employs general-purpose LLMs in a zero-shot setting to reason directly over extracted ion chromatograms and generate human-interpretable rationales for each decision. On an expert-annotated *Streptococcus pyogenes* DIA benchmark dataset, ChatDIA achieved 96.9% accuracy, matching the domain-specific state-of-the-art software DIA-NN (95.5%). In a challenging single-cell HEK-293T DIA proteomics dataset, ChatDIA further demonstrated excellent performance, achieving a lower risk-coverage area under the curve than DIA-NN (0.06 vs. 0.12) and identifying 38.75% and 46.25% of library peptides at 1% and 5% false discovery rate, respectively, compared with 16.25% and 48% for DIA-NN. Together, these results demonstrate that zero-shot LLM reasoning can competitively automate core targeted DIA decision-making while providing transparent, inspectable rationales that enable conversational, interactive validation and data exploration in noisy proteomics applications. More broadly, ChatDIA illustrates how reasoning-based AI systems can move beyond prediction to generate evidence-based decisions, offering a foundation for more general and interpretable computational analysis in proteomics.

## Introduction

Virtually all cellular processes are carried out by proteins. The complete set of proteins expressed in a cell, tissue, or bodily fluid at a given time, collectively termed the proteome, provides a functional link between the genome and biological processes and phenotypes (1). Proteomics aims to comprehensively characterize the proteome and has become increasingly important for understanding cellular behavior, identifying therapeutic targets, and advancing personalized medicine (2–4). Liquid chromatography coupled with tandem mass spectrometry (LC–MS/MS) is a central technology for large-scale proteomics (2). Over the past decade, data-independent acquisition (DIA) has emerged as a widely adopted strategy for high-throughput, reproducible, and large-scale proteome profiling (5–7). DIA systematically fragments all peptide precursors within predefined mass-to-charge (m/z) isolation windows, enabling consistent peptide detection and quantification across samples and experimental runs (8–10).

Despite these advantages, DIA data analysis remains inherently challenging due to the highly multiplexed nature of MS/MS spectra, which contain fragment ions from multiple co-fragmented precursors. To address this complexity, DIA analysis is typically performed using peptide-centric, library-based approaches (5, 11), in which prior information about peptide fragmentation and retention time is compiled into a spectral library and queried against the DIA data. Chromatograms corresponding to precursor and fragment ions of interest are then extracted, and the central analytical challenge is to identify the correct chromatographic peak group among competing alternatives and interferences (11, 12).

A range of tools has been developed to address this task, spanning manual inspection, classical machine learning models, and deep neural networks. Visualization tools such as Skyline (13) enable expert users to manually inspect extracted ion chromatograms (XICs), often achieving high accuracy; however, manual validation is labor-intensive, subjective, and infeasible for large-scale or high-throughput studies (14). Algorithmic approaches instead compute various features for candidate peak groups, such as peak shape, fragment-ion co-elution, and intensity consistency, which serve as the basis for downstream scoring and discrimination. These features may be explicitly defined, as in OpenSWATH (12), Spectronaut (9), and DIA-NN (15), or implicitly learned by deep learning models such as DeepMRM (16), DreamDIA (17), and DIA-BERT (18), which are trained on labeled mass spectrometry data. Based on these extracted features and learned model parameters, initial peak group-level scores or probabilities are generated. In many workflows, these scores are further refined using additional statistical or machine learning frameworks (19), most commonly target-decoy strategies combined with semi-supervised discriminant modeling (14, 20–22), to produce final discriminative scores and to estimate false discovery rate (FDR). However, the decision-making processes of existing DIA analysis tools are often opaque and difficult to interrogate. Composite discriminant scores derived from numerous features provide limited insight into why a specific peak group is selected over competing alternatives. These limitations are particularly pronounced in challenging applications such as single-cell proteomics, which is characterized by low signal-to-noise ratios and substantial spectral interference (23), where manual inspection is frequently required to validate automated results.

Recent advances in large language models (LLMs) have demonstrated that general-purpose models, typically trained on large-scale text corpora, can perform complex tasks without task-specific training (24). Modern LLMs are typically based on transformer architectures, which use self-attention mechanisms to model long-range dependencies in sequential data (25). Scaling these models to large and diverse training datasets has led to emergent capabilities in zero-shot and few-shot learning, enabling generalization to previously unseen tasks (26–28). Beyond language understanding, LLMs have been shown to perform multi-step reasoning when prompted appropriately (29). Techniques such as chain-of-thought (29) prompting explicitly encourage models to articulate intermediate reasoning steps, improving performance on arithmetic, logical, and symbolic reasoning tasks, including in zero-shot settings without labeled training data (30).

To date, applications of pre-trained LLMs in scientific domains have largely focused on text-centric tasks such as literature mining, code generation, and hypothesis summarization. Their potential role in core analytical decision-making, particularly within quantitative data analysis pipelines, remains largely unexplored. In proteomics, existing deep learning approaches are typically task-specific models trained directly on mass spectrometry data and optimized for classification or regression objectives. Whether a general-purpose LLM, without any domain-specific training on mass spectrometry data, can perform essential analytical tasks such as chromatographic peak group evaluation and true signal selection has not been systematically investigated.

Here, we introduce ChatDIA, a large language model-based workflow that performs targeted DIA analysis through an explicit reasoning-based decision framework. ChatDIA operates in two complementary modes: an API-based mode for fully automated peptide-level decision-making, and an interactive mode that enables natural language-based interaction for human-in-the-loop inspection, validation, and data exploration. Rather than relying on specialized, domain-trained models, ChatDIA employs general-purpose LLMs in a zero-shot setting to reason directly over extracted chromatographic signals and to generate human-interpretable rationales for each peptide-level decision. When evaluated on an expert-annotated *Streptococcus pyogenes* DIA proteomics benchmark comprising 423 randomly selected precursors, ChatDIA showed high agreement with manual annotations, achieving 96.9% accuracy and modestly outperforming DIA-NN (95.5%). In the context of a low signal-to-noise single-cell HEK-293T DIA dataset containing 400 randomly selected precursors (244 with true peptide signal and 156 without by manual annotation), ChatDIA demonstrated favorable risk-coverage performance compared to DIA-NN, as reflected by a lower area under the curve (0.06 vs. 0.12). Across commonly used FDR thresholds, ChatDIA identified 38.75% and 46.25% of peptides at 1% and 5% FDR, respectively, compared with 16.25% and 48% achieved by DIA-NN. Beyond competitive analytical performance, ChatDIA uniquely enables conversational interaction with the analysis process, providing inspectable, case-specific rationales that support transparent decision-making and facilitate validation in noisy, interference-dominated data regimes.

## Material and Methods

### Benchmarking datasets

To benchmark and evaluate the performance of ChatDIA, we used two previously published proteomics datasets representing bulk and single-cell DIA experiments.

The first dataset (12, 31) consists of 16 SWATH DIA runs acquired from the *Streptococcus pyogenes* bacterial strain SF370 and is available via PeptideAtlas (PASS00788). Data was acquired on a SCIEX 5600 TripleTOF instrument. For this dataset, we leveraged an existing assay library, OpenSWATH (v2.4.0) (12) results coupled with PyProphet (v2.1.3) (14, 21) FDR correction, and a set of 437 randomly selected, manually annotated precursors from a previously published study (32, 33) (PeptideAtlas PASS01508). From this dataset, we used four technical replicates corresponding to *S. pyogenes* samples grown in 0% human plasma. A total of 423 precursors (transition groups) that had manual annotations in at least one of the four replicates and corresponding OpenSWATH results were retained for downstream analysis. To ensure annotation accuracy, all selected precursors were subjected to additional manual validation. For comparative analysis, DIA-NN (v1.8) was run on the full *S. pyogenes* dataset as described previously (34).

The second dataset (23) is a single-cell HEK-293T proteomics dataset available from the MassIVE database (MSV000093867), acquired on a Bruker timsTOF SCP instrument. One run from this dataset was selected and processed using OpenSWATH (v3.5.0) with PyProphet (v3.0.5) for FDR estimation, as well as DIA-NN (v2.2.0 Academia) for comparative analysis. Based on the OpenSWATH results, 400 unique precursors (transition groups) were chosen: 200 precursors were randomly selected from identifications with an FDR below 1%, and 200 were randomly selected from those above this threshold. Manual annotation identified true peptide signals in 244 of these precursors, while the remaining 156 showed no evidence of target peptide signal.

### DIA data processing and chromatogram extraction

DIA ion chromatograms were extracted, picked and scored by OpenSWATH using the default parameters. For each transition group, OpenSWATH-generated extracted ion chromatograms (XICs) were further processed by the ChatDIA workflow into three alternative data representations: numerical retention time-intensity pair arrays, XIC images, and ion trace images (Supplementary Figure 1).

ChatDIA was operated in two different settings. In the first setting, the model was provided only with raw extracted ion chromatograms (XICs), together with assay library metadata as reference information, for direct peptide identification. In the second setting, the model was provided with one to five candidate peak group positions per chromatogram, including their retention times and, when available, intensities, and was tasked with selecting the correct peak group from among the proposed candidates. Candidate peak groups could be proposed either by simple heuristic algorithms or by an LLM acting as a candidate proposer within the ChatDIA workflow. No OpenSWATH-derived peak group scores, rankings, or confidence estimates were supplied; thus, ChatDIA had no access to OpenSWATH-calculated scoring metrics.

### ChatDIA model configuration and prompting

ChatDIA was implemented using OpenAI large language models GPT-5-mini, GPT-5.1, GPT-5.2, and GPT-5.4. The automated version of ChatDIA utilizes OpenAI API batch processing for large-scale analysis, while a lightweight interactive version is deployed within OpenAI ChatGPT. All models were run with reasoning (“thinking”) mode enabled and set to the “high” level.

Prompts provided to the models included a high-level task description for peptide signal identification, the selected XIC data representation, relevant assay library metadata, either with or without structured guidance to support multi-criteria evaluation of chromatographic evidence. The prompt without explicit guidance on peak detection and evaluation served as a baseline, representing model performance without expert knowledge explicitly encoded in the prompt. Example prompts and corresponding model outputs are shown in Supplementary Figure 1.

### Evaluation metrics and statistical analysis

For the *Streptococcus pyogenes* dataset, where most annotated precursors were expected to be present, peak group selection performance was primarily evaluated using accuracy.

For both the *Streptococcus pyogenes* and single-cell HEK-293T datasets, confidence-aware decision reliability was assessed using risk-coverage (FDR-coverage) analysis. Manual validation was treated as ground truth for decision correctness: a positive label indicates the model’s decision (selected peak RT boundaries or no peak) matches manual inspection. Coverage was defined as the fraction of predictions at or above a given confidence threshold among all manually annotated transition groups; mathematically, coverage = (TP + FP) / (TP + FP + TN + FN). The confidence threshold was determined using method-specific confidence measures: LLM peptide identification confidence scores or ensemble-derived scores from multiple independent LLM runs for ChatDIA, 1 − q values for DIA-NN, and discriminative scores for OpenSWATH. The FDR was defined as the fraction of retained predictions that were incorrect by manual validation, FP / (TP + FP). An example of model score thresholding, coverage estimation, and FDR calculation is provided in Supplementary Table 1.

For the single-cell dataset, standard precision-recall analysis was additionally performed to evaluate model performance.

## Results

### ChatDIA enables automated DIA peptide identification with natural language-based interaction

We developed ChatDIA, a zero-shot large language model (LLM)-based workflow for targeted DIA proteomics analysis (Figure 1). ChatDIA integrates experimental library metadata with DIA XICs and delegates peak group detection, when peak positions are not precomputed by simple algorithms or an LLM, and peak group selection to an LLM. For each transition group, the LLM evaluates multiple sources of chromatographic evidence, including peak shape, fragment-ion co-elution, potential interference and similarity of peptide properties to the library (relative fragment ion intensity, retention time) to identify the most plausible peptide signal. The model produces a final confidence score and provides a natural-language justification for its decision. Examples of the model’s evaluations are shown in Supplementary Figure 1. This formulation enables peak group selection to be performed as an explicit, multi-criteria reasoning process rather than as an opaque optimization over learned scoring functions.

**Figure 1.**
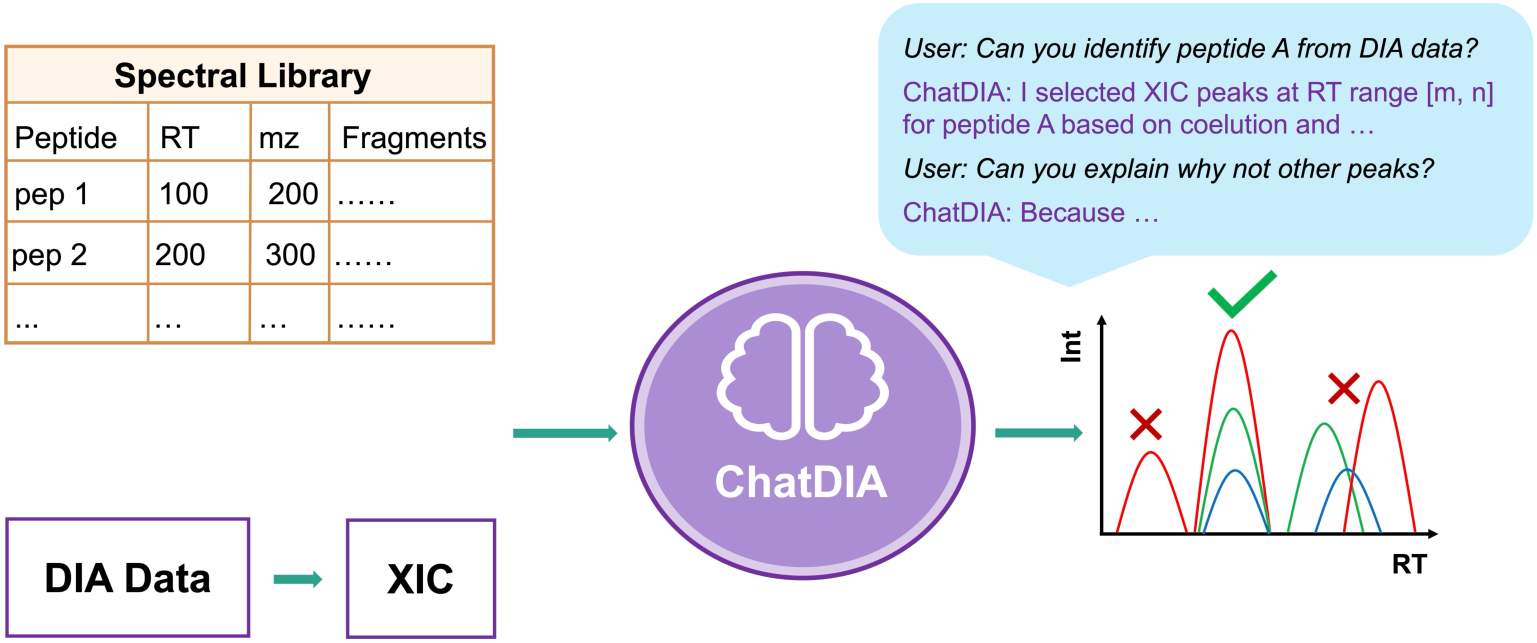
Overview of ChatDIA, a large language model workflow for DIA proteomics. ChatDIA integrates an experimental library with DIA extracted ion chromatograms (XICs) generated by external software (e.g., OpenSWATH) and delegates peak group detection (when not precomputed) and peak selection to a large language model (LLM). For each transition group, the LLM evaluates chromatographic evidence and assay library information to identify the peak group corresponding to the target peptide, while generating human-readable rationales to facilitate interactive review. ChatDIA supports both automated analysis for large-scale studies and natural language-based interaction for peak selection, validation, error diagnosis, and exploratory data analysis.

ChatDIA operates in two complementary modes. In its primary mode, API-based LLM inference enables fully automated analysis, producing peptide identification decisions together with confidence estimates and human-readable explanations suitable for downstream filtering and quality control. In parallel, ChatDIA supports interactive usage through web-based or application-level interfaces that expose the same reasoning capabilities in a conversational format. This interactive mode places particular emphasis on transparent, human-readable explanations, allowing users to inspect individual peak selection decisions, query model rationales, diagnose errors, and explore chromatographic evidence using natural language, thereby bridging automated large-scale DIA analysis with human-in-the-loop data interpretation.

### Numerical chromatogram representations outperform image-based inputs for LLM peak selection

We first evaluated how different representations of DIA XICs influence LLM performance in peak group selection. Using 423 unique transition groups measured across four technical replicates (1,603 total evaluations) from a *Streptococcus pyogenes* proteomics dataset, we assessed ChatDIA accuracy across three input modalities: numerical retention time-intensity arrays, XIC images, and ion trace image representations (Figure 2). Examples of prompts, XIC representations, and corresponding model inputs and outputs are shown in Supplementary Figure 1. For numerical array and XIC image inputs, model performance was evaluated under two different configurations: (1) providing only raw XIC data and assay library metadata, and (2) providing candidate peak group positions (retention times and intensities), from which the model was tasked to select the correct one. Ion trace image inputs were evaluated with candidate peak group information embedded directly within the prompt.

**Figure 2.**
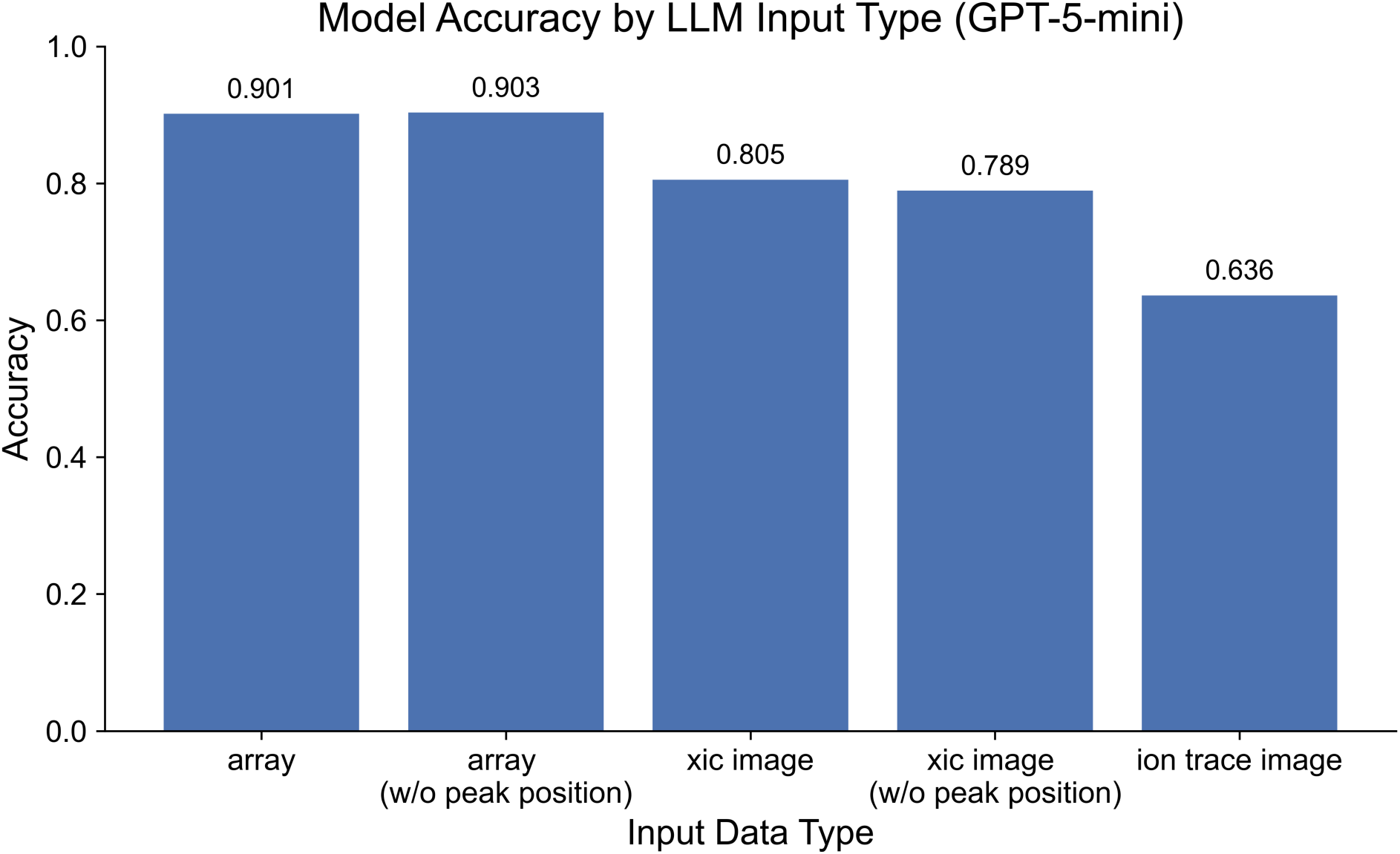
Model accuracy across different LLM input data representations. Accuracy of ChatDIA (using OpenAI GPT-5-mini) in identifying the correct peak group on the *Streptococcus pyogenes* proteomics dataset, evaluated using 423 unique transition groups measured across four technical replicates (1,603 total evaluations). Model accuracy is compared across multiple representations of extracted ion chromatograms (XICs), including numerical retention time-intensity arrays, XIC images, and ion trace images. For numerical array and XIC image inputs, performance is assessed both with OpenSWATH-provided peak position information (retention time start/stop and intensity; denoted as “array” and “xic image”) and without this information (denoted as “array (w/o peak position)” and “xic image (w/o peak position)”). Ion trace image (“ion trace image”) inputs are evaluated with candidate peak group information embedded in the LLM input.

In all experiments, OpenSWATH was used exclusively for chromatogram extraction only (mode 1) or for chromatogram extraction combined with peak group candidate proposal, including retention-time boundaries and intensity estimates for candidate peak groups (mode 2). No peak group scores, rankings, or confidence estimates from OpenSWATH were provided to the LLM. As a result, peak selection decisions were driven solely by the model’s reasoning over the supplied chromatographic evidence and assay library metadata, rather than by conventional DIA scoring functions.

ChatDIA (using OpenAI GPT-5-mini) achieved the highest accuracy when XICs were provided as numerical retention time-intensity arrays, with consistently strong performance (accuracy) both with (0.901) and without (0.903) candidate peak group, suggesting that the model is capable of selecting peak groups in raw chromatographic data unassisted by external tools. In contrast, XIC image representations yielded lower accuracy (0.805 with peak positions and 0.789 without), while ion trace image inputs resulted in substantially reduced performance (0.636). Together, these results demonstrate that explicit numerical representations of chromatographic data more effectively support zero-shot, multi-criteria reasoning by LLMs than image-based encodings.

### ChatDIA matches state-of-the-art DIA tools on a curated *Streptococcus pyogenes* proteomics dataset without task-specific training

We next compared ChatDIA against established DIA analysis tools using the same *Streptococcus pyogenes* proteomics dataset with expert-annotated ground truth. Performance was evaluated across 423 unique precursors measured in four technical replicates, yielding 1,603 chromatograms with 4,783 candidate peak groups and 281,834 chromatographic positions to select from. ChatDIA (GPT-5.1 and GPT-5.2) was compared against DIA-NN and OpenSWATH (Figures 3 and 4). For ChatDIA, DIA XICs were provided as numerical retention time-intensity arrays and evaluated both with and without precomputed candidate peak group retention times and intensities.

**Figure 3.**
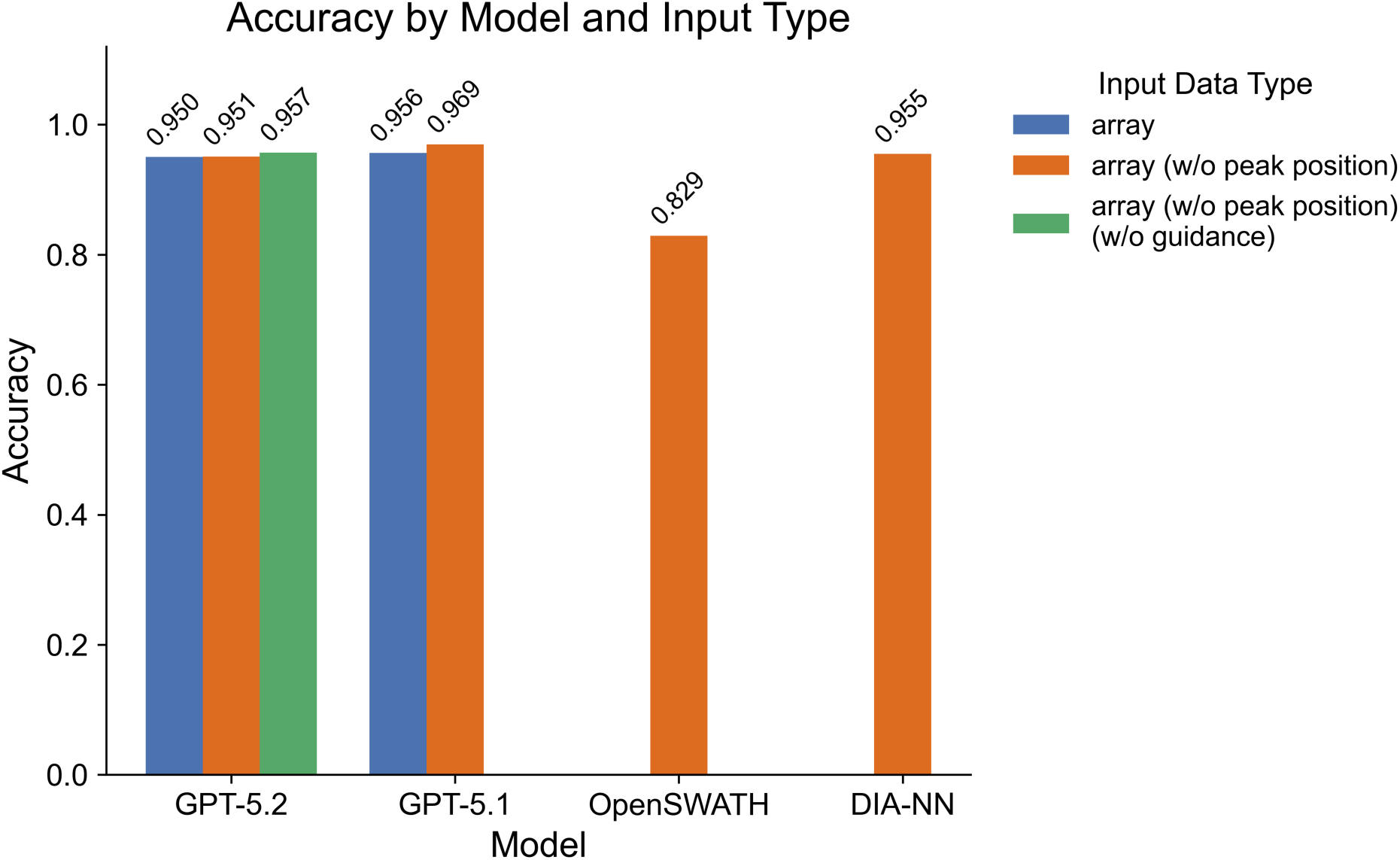
Comparison of model and algorithm accuracy on the *Streptococcus pyogenes* proteomics dataset. Performance of ChatDIA with GPT-5.1, ChatDIA with GPT-5.2, DIA-NN, and OpenSWATH was evaluated using 423 unique transition groups measured across four technical replicates (1,548 total evaluations for DIA-NN due to missing peptide detections; 1,603 evaluations for all other methods). XICs were provided as numerical retention time-intensity arrays. Orange bars denote inputs without additional peak position information, blue bars denote inputs augmented with candidate peak positions (retention time start/end and intensities), and green bars denote ChatDIA performance without guidance (no explicit knowledge of how to detect, evaluate, or score peaks was provided) in addition to the absence of peak position. OpenSWATH and DIA-NN results were computed without FDR filtering.

**Figure 4.**
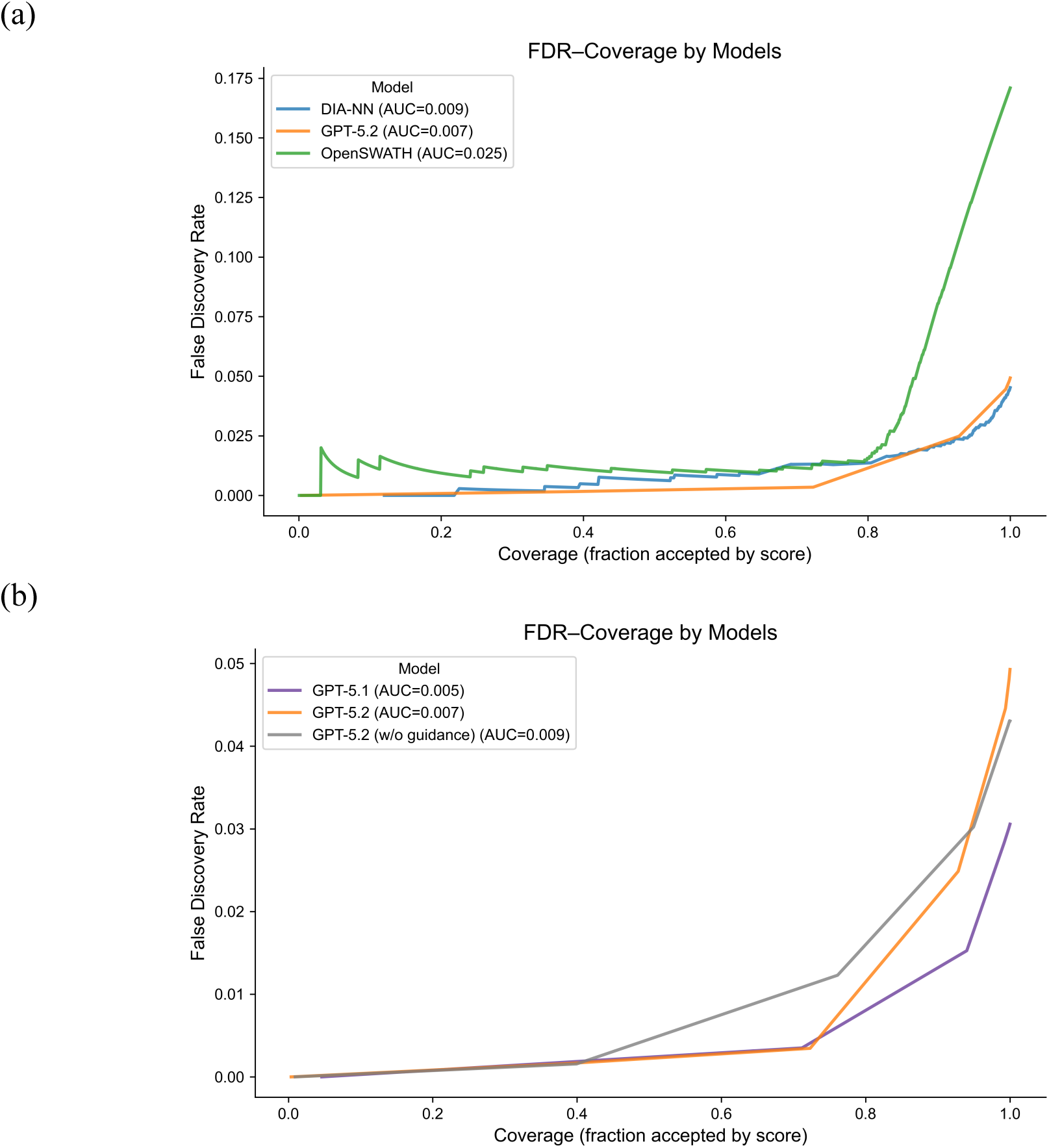
Risk-coverage analysis of top-1 peak group selection on the *Streptococcus pyogenes* proteomics dataset. Risk-coverage (FDR-coverage) analysis comparing ChatDIA with established DIA analysis algorithms. **(a)** FDR-coverage curves for ChatDIA with GPT-5.2 using numerical retention time-intensity array inputs without peak information, DIA-NN, and OpenSWATH. Performance was evaluated using 423 unique transition groups measured across four technical replicates (1,548 total evaluations for DIA-NN due to missing peptide detections; 1,603 evaluations for the other methods). Each point on the curve is obtained by varying a method-specific confidence threshold (LLM confidence scores for ChatDIA, FDR-adjusted discriminative scores for OpenSWATH, and 1−q values for DIA-NN). At each threshold, coverage is defined as the fraction of model predictions with confidence scores greater than or equal to the threshold; mathematically, coverage = (TP + FP) / (TP + FP + TN + FN). FDR is defined as the fraction of retained top-1 peak group selections rejected by manual validation, FP / (TP + FP). **(b)** FDR-coverage curves comparing ChatDIA configurations using numerical XIC array inputs under different guidance settings, including GPT-5.2 with explicit guidance, GPT-5.1 with explicit guidance, and GPT-5.2 without explicit guidance.

Using numerical array inputs, ChatDIA achieved peak group selection accuracy comparable to state-of-the-art DIA tools (Figure 3). In the baseline setting, where no guidance on chromatogram or peak evaluation was provided in the prompt and no candidate peak positions were supplied, ChatDIA with GPT-5.2 achieved an accuracy of 0.957. When explicit guidance for peak detection and evaluation was included, ChatDIA with GPT-5.2 reached accuracies of 0.950 and 0.951 with and without candidate peak group positions, respectively, while ChatDIA with GPT-5.1 achieved accuracies of 0.956 and 0.969 under the same conditions. These results are comparable to DIA-NN (0.955) and substantially higher than OpenSWATH (0.829). Notably, ChatDIA achieved this level of performance using zero-shot LLM inference, without any mass spectrometry-specific model training or task-specific parameter optimization.

Beyond overall accuracy, we evaluated the reliability of top-1 peak group selection using FDR-coverage analysis. FDR-coverage curves were generated for ChatDIA with GPT-5.2 and GPT-5.1 without candidate peak group information, for ChatDIA with GPT-5.2 in the absence of both prompt guidance and candidate peak group information, and for DIA-NN and OpenSWATH on the same dataset (Figure 4). This analysis assesses, across varying FDR thresholds, the number of peptides identified by each method. Overall performance was summarized using the area under the risk-coverage curve (AUC), with lower values indicating better performance.

With explicit prompt guidance emphasizing retention-time proximity to the spectral library, coelution, peak shape, and fragment-ion patterns, ChatDIA with GPT-5.2 exhibited strong risk-coverage behavior, achieving a substantially lower AUC (0.007) than both DIA-NN (0.009) and OpenSWATH (0.025). At an FDR threshold of 1%, ChatDIA with GPT-5.2 achieved 72.3% coverage, exceeding that of DIA-NN and OpenSWATH (64.7% each). ChatDIA with GPT-5.1 showed comparable performance, with an AUC of 0.005 and 71.2% coverage at 1% FDR. In contrast, in the baseline setting without explicit prompt guidance, ChatDIA with GPT-5.2 exhibited reduced confidence calibration, with an AUC of 0.009 and only 39.9% coverage at 1% FDR.

To assess whether prior public availability of the *Streptococcus pyogenes* proteomics dataset and potential exposure during GPT pretraining influenced model performance, we masked peptide identities and applied a +200 s shift to retention times (including library RTs), and evaluated ChatDIA with GPT-5.2 using numerical XIC array inputs without peak position information (Supplementary Figure 2). Performance remained comparable under these perturbations, with identical accuracy (95.1%), similar risk-coverage behavior (AUC = 0.009 for anonymized inputs versus 0.007 for original inputs), and nearly identical peptide identification rates at 1% FDR (72.2% vs. 72.3%). These results suggest that model performance is driven by data-derived reasoning rather than memorization of prior information.

Together, these results demonstrate that confidence estimates derived from LLM-based, multi-criteria reasoning over numerical chromatographic data provide a robust basis for selectively accepting high-confidence peptide identifications. They further indicate that, while overall peak selection accuracy can remain high, explicit expert guidance in prompt design substantially improves the reliability and calibration of model-derived confidence scores, underscoring the value of domain knowledge in instructing LLMs for proteomics data analysis.

### ChatDIA enables robust peptide identification in single-cell proteomics through abstention-aware modeling

ChatDIA was further evaluated on a challenging single-cell HEK-293T proteomics DIA dataset characterized by low signal-to-noise ratios and substantial spectral interference. A total of 400 precursors were randomly selected for manual annotation and model evaluation. The performance of ChatDIA was compared against DIA-NN and OpenSWATH using risk-coverage (FDR-coverage) and precision-recall analyses (Figure 5). Confidence thresholds were determined using method-specific confidence measures: LLM confidence scores or ensemble-derived scores from multiple independent LLM runs for ChatDIA, 1 − q values for DIA-NN, and MS2-level discriminative scores for OpenSWATH.

**Figure 5.**
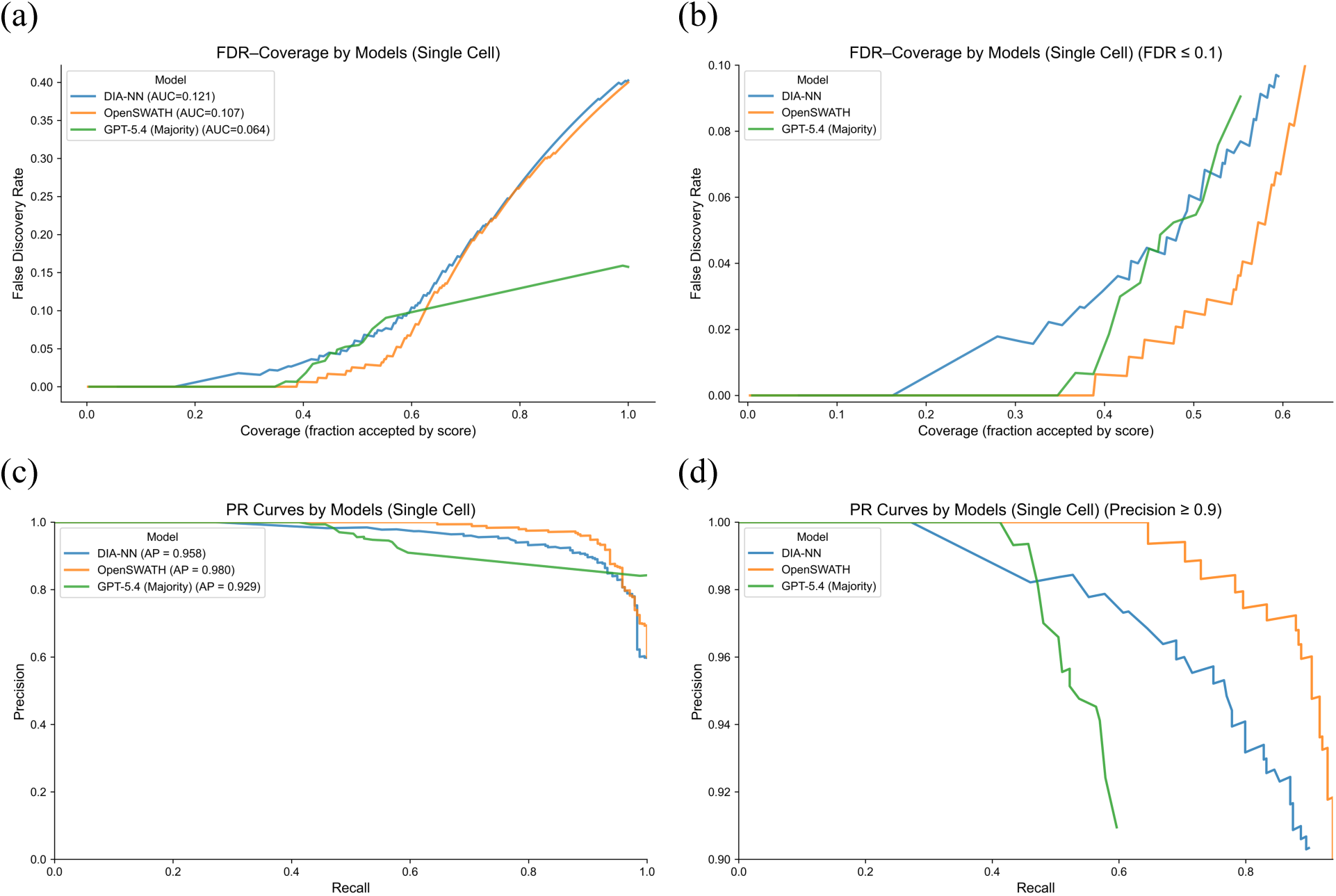
Model performance comparison on a single-cell proteomics dataset. **(a-b)** Risk-coverage analysis and **(c-d)** precision-recall (PR) analysis comparing ChatDIA with GPT-5.4 (ensemble of three runs with majority voting), DIA-NN, and OpenSWATH on a single-cell HEK-293T proteomics dataset. ChatDIA models use numerical XIC array representations incorporating candidate peak group positions, with Gaussian smoothing (σ = 1.0), and allow abstention (i.e., reporting no peak group). A total of 400 transition groups were selected for manual annotation based on OpenSWATH results, comprising 200 peptides with identifications below an FDR threshold of 0.01 and 200 above this threshold. Performance was assessed using risk-coverage analysis. Coverage is defined as the fraction of predictions with confidence scores greater than or equal to a given score threshold among the 400 manually annotated transition groups (mathematically, coverage = (TP + FP) / (TP + FP + TN + FN)), where the threshold is determined by method-specific confidence measures (LLM confidence scores for ChatDIA, 1−q values for DIA-NN, and discriminative scores for OpenSWATH). FDR is defined as the fraction of retained predictions rejected by manual validation, FP/(TP+FP). Precision-recall analysis was performed using standard PR metrics. Panels (a) and (c) show risk-coverage and PR curves over the full operating range, respectively, whereas panels (b) and (d) show zoomed views focusing on FDR ≤ 0.1 and precision ≥ 0.9, respectively.

We first evaluated the baseline performance of ChatDIA using GPT-5.2 without explicit prompt guidance for peak evaluation and without candidate peak group information. Numerical retention time-intensity arrays were used as input, and Gaussian smoothing was applied (Gaussian kernel standard deviation σ = 1 scan, corresponding to approximately 0.74 s on the sampled retention-time grid). Under these conditions, ChatDIA demonstrated poor confidence calibration, achieving an area under the FDR-coverage curve (AUC) of 0.225 (Supplementary Figure 3). No peptides were confidently identified within 5% FDR, indicating that the LLM was unable to reliably resolve peptide signals in highly noisy, interference-rich single-cell data in the absence of additional structural constraints or task-specific guidance.

Given the low signal-to-noise characteristics of the dataset, we systematically explored prompt design strategies to improve robustness, including the provision of candidate peak group positions to reduce the search space and varying degrees of chromatogram smoothing to suppress noise. In addition, to address the model’s tendency toward overconfident predictions, we explicitly permitted ChatDIA to abstain from reporting a peak group when model confidence was insufficient. The effects of candidate peak group information, abstention behavior, and chromatogram smoothing strength were evaluated using FDR-coverage analysis with GPT-5.2 (Supplementary Figure 3). Providing candidate peak group positions consistently improved performance, while enabling abstention further enhanced confidence calibration and risk-coverage behavior. Moderate Gaussian smoothing (σ = 1) yielded overall best performance, whereas both unsmoothed raw chromatograms and excessive smoothing degraded coverage and confidence calibration. While LLM with raw data yielded higher coverage at 1% FDR (Supplementary Figure 3b, Supplementary Table 2), the sharply increasing risk at higher coverage indicate that it may not be the optimal choice overall.

Building on the initial GPT-5.2 evaluation, we next assessed ChatDIA using a newer LLM, GPT-5.4. Under this setting, ChatDIA was provided with OpenSWATH-precomputed candidate peak group retention times and intensities, moderate Gaussian smoothing was applied (σ = 1), and the model was explicitly permitted to abstain. Performance was evaluated across three independent runs and different ensemble strategies (Figure 5 and Supplementary Figure 4). Under these conditions, ChatDIA with GPT-5.4 using majority voting achieved the most favorable overall risk-coverage performance, attaining the lowest AUC (0.064), outperforming DIA-NN (0.121) and OpenSWATH (0.107). Notably, in the very low FDR regime, OpenSWATH demonstrated the strongest performance. At a stringent FDR threshold of 1%, ChatDIA with GPT-5.4 (majority voting) achieved peptide identification coverage of 38.75%, exceeding DIA-NN (16.25%) but remaining slightly lower than OpenSWATH (42.5%). Detailed identification rates at FDR thresholds of 1%, 5%, and 10% are provided in Supplementary Table 2.

Individual GPT-5.4 runs showed moderate variability (Supplementary Figure 4 and Supplementary Table 2), but consistently outperformed DIA-NN at FDR ≤ 0.01. Among ensemble strategies, majority voting provided improved performance in low-FDR regions compared to conservative and open strategies (Supplementary Figure 4 and Supplementary Table 2). Detailed definitions of ensemble strategies are provided in Supplementary Table 3.

Precision-recall analysis (Figure 5c, d) showed that ChatDIA achieved slightly lower discrimination performance than DIA-NN and OpenSWATH. Average precision values were 0.98 for OpenSWATH, 0.958 for DIA-NN, and 0.929 for ChatDIA with GPT-5.4. These results indicate that ChatDIA is highly competitive and performs comparably to conventional tools under standard precision-recall metrics.

Together, these results indicate that ChatDIA’s LLM-based, abstention-aware reasoning framework provides a principled approach for mitigating overconfident predictions in single-cell proteomics. By explicitly allowing abstention and leveraging multi-criteria reasoning over numerical chromatographic data, ChatDIA supports reliable peptide identification under low-signal, interference-dominated single-cell DIA conditions.

### Multi-agent ChatDIA supports direct peak group detection and selection in single-cell proteomics

To assess whether ChatDIA with GPT-5.4 could operate robustly without OpenSWATH-proposed candidate peak groups (including their retention times and intensities), we evaluated two ChatDIA workflow configurations (Supplementary Figure 5). In the first configuration, a multi-agent workflow was used, in which one LLM proposed candidate peak groups and a second LLM selected the most likely true peptide signal from the proposed candidates. In the second configuration, a single LLM was tasked with both peak group detection and selection. Numerical XIC arrays with Gaussian smoothing (σ = 1) were provided to all agents, and the model was explicitly allowed to abstain when selecting true peak groups. Three independent ChatDIA runs, together with their ensemble outputs using the same strategies described above, were evaluated by FDR-coverage analysis (Supplementary Figure 6 and Supplementary Table 4). The multi-agent ChatDIA workflow showed better overall performance than the single-agent setting. At 1% FDR, multi-agent ChatDIA identified 25.50% of precursors using either majority voting or conservative voting ensembles, whereas the three individual runs achieved precursor identification coverages of 15.75%, 7.00%, and 17.25%. In contrast, single-agent ChatDIA identified 14.25% of precursors using the conservative voting ensemble and 3.75% using majority voting, with individual run coverages of 0%, 3.75%, and 16.25% at 1% FDR. These results suggest that separating candidate peak group proposal and peak group selection across two LLM agents improves performance compared with assigning both tasks to a single LLM. Although performance was lower than that achieved when OpenSWATH-proposed candidate peak groups were provided to the LLM, the multi-agent workflow still achieved performance comparable to DIA-NN. These results indicate that ChatDIA can support direct peak group detection and selection without relying exclusively on externally precomputed candidate peak groups.

### ChatDIA provides interpretable results and enables natural language-based interaction

In addition to automated analysis, ChatDIA is designed to support interpretable decision-making and interactive exploration of DIA data. Unlike conventional DIA tools that output fixed scores or ranked peak groups, ChatDIA produces explicit natural-language explanations that summarize the chromatographic evidence and reasoning underlying each peak selection decision.

ChatDIA supports interactive use through a lightweight application implemented within OpenAI ChatGPT, exposing the same reasoning capabilities used in large-scale analyses in a conversational interface (Figure 6a). In this mode, users can interact with the system via natural language to perform DIA-related tasks, such as peptide signal identification, XIC visualization, explanation of the rationale underlying individual peak group selections, and literature searches for proteins of interest (Supplementary Figure 7). This interaction paradigm enables users to inspect chromatographic evidence, validate model decisions, and explore ambiguous cases in a human-in-the-loop manner, complementing automated large-scale workflows.

**Figure 6.**
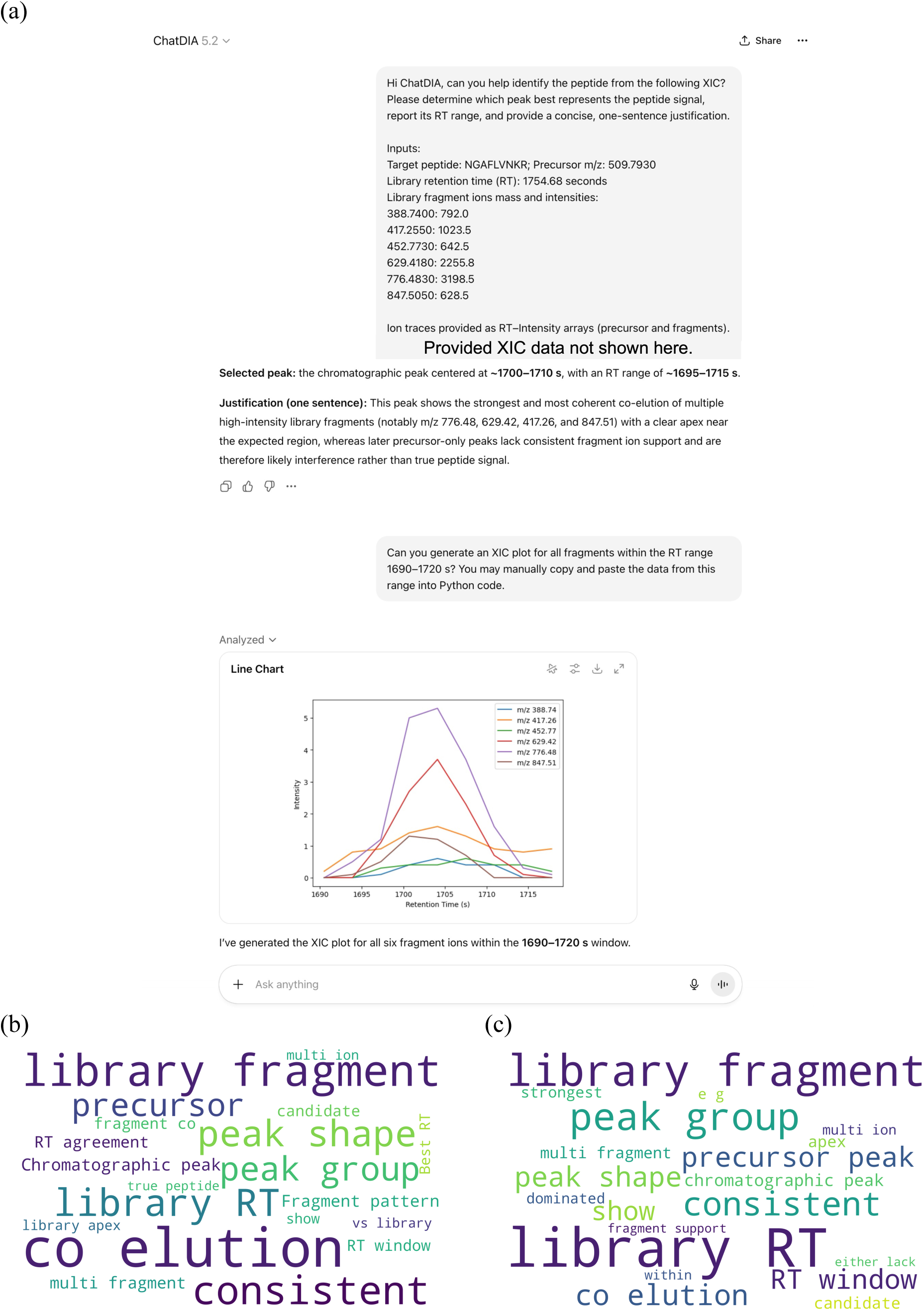
Unique interactive and interpretable features of ChatDIA. **(a)** Example of the ChatDIA application in a lightweight mode implemented within OpenAI ChatGPT, illustrating interactive user engagement for DIA data analysis. The example shows that users can directly query the model using natural language to perform DIA analyses, such as identifying peptide signals and generating diagnostic plots. The screenshot is taken from the OpenAI ChatGPT platform, on which the current lightweight version of ChatDIA is deployed. **(b-c)** Word cloud analysis of LLM-generated decision justifications. The visualizations summarize justification text produced by ChatDIA with GPT-5.2 on the *Streptococcus pyogenes* proteomics dataset using numerical retention time-intensity array inputs without peak positions provided. Word size reflects the relative frequency of terms appearing in the model-generated justifications. Panel (b) corresponds to ChatDIA with explicit guidance, and panel (c) corresponds to ChatDIA without explicit guidance.

To characterize the interpretability of ChatDIA’s decision process, we analyzed the natural-language justifications generated by the model during peak group selection. Word clouds summarizing decision justifications produced by ChatDIA with GPT-5.2 on the *Streptococcus pyogenes* proteomics dataset are shown in Figure 6b and c. The most frequently occurring terms reflect core chromatographic evaluation criteria, including fragment-ion co-elution, peak shape consistency, and retention time agreement to the library. The prevalence of these terms indicates that the model’s explanations consistently reference established principles used by expert analysts when evaluating DIA chromatograms, rather than relying on opaque or uninterpretable heuristics. In addition, dozens of ChatDIA outputs were reviewed by multiple human experts and found to be generally reasonable.

Together, these results demonstrate that ChatDIA not only achieves competitive performance in automated DIA analysis but also provides transparent, human-readable rationales and supports natural language-based interaction. This combination enables both scalable automated peptide identification and interactive, expert-guided data interpretation, bridging traditional DIA workflows with emerging LLM-driven analytical paradigms.

## Discussion

In this work, we introduce ChatDIA, a zero-shot LLM-based framework for DIA proteomics that integrates automated analysis with interpretable, natural language-driven reasoning. Unlike conventional purpose-built DIA tools that rely on mass spectrometry-specific training or fixed algorithmic scoring functions, ChatDIA applies zero-shot reasoning directly to explicit chromatographic evidence to perform peak group selection. Our results demonstrate that this LLM-based approach can achieve performance comparable to state-of-the-art DIA methods while providing transparency and interactive capabilities through natural language that are not available in existing DIA workflows.

The primary novelty of this work lies in the use of general-purpose LLMs, without mass spectrometry-specific training or fine-tuning, as reasoning engines operating over structured experimental data. More broadly, this represents a paradigm shift from specialized, task-specific algorithms toward general-purpose models that derive decisions through structured reasoning over experimental evidence rather than learned scoring functions. Traditional proteomics machine learning methods depend on curated training datasets and are often tightly coupled to specific acquisition schemes, instruments, or experimental conditions (9, 12, 14–19, 21, 22). In contrast, ChatDIA operates in a zero-shot setting, where structured prompts define the analytical objective and provide relevant chromatographic evidence, enabling the model to reason directly over the data.

In practice, this reasoning-based approach operates by evaluating competing interpretations of chromatographic evidence. DIA peak group selection inherently requires assessing relational patterns, including chromatographic peak shape, fragment-ion co-elution, and consistency with assay libraries. These criteria are qualitative, multi-dimensional, and routinely applied by human experts during manual validation. ChatDIA formalizes this process by enabling the model to construct and compare candidate explanations, selecting the interpretation that best accounts for the observed signal. As a result, the model output represents not only a prediction but a data-supported decision grounded in explicit reasoning over chromatographic structure.

A key advantage of the ChatDIA framework is that it does not impose a tradeoff between automation and interpretability. Through API-driven execution, ChatDIA enables scalable analysis across large datasets, producing peak selection decisions accompanied by confidence estimates and human-readable rationales. These outputs support downstream filtering, quality control, and error analysis. At the same time, the same reasoning process can be accessed interactively, allowing users to inspect individual cases, examine chromatographic evidence, and interrogate model decisions through natural language. This dual-mode design bridges automated analysis and human-in-the-loop interpretation, which is particularly valuable in challenging settings such as single-cell proteomics, where signal quality is limited and uncertainty is intrinsic.

An interesting observation from this study is that, in complex datasets such as single-cell proteomics, LLM-based inference exhibited greater run-to-run variability than deterministic algorithms. This behavior may resemble manual annotation, where expert judgments can vary across repeated evaluations of ambiguous cases. Although such variability may appear to be a limitation, it also provides diversity that can be leveraged through ensemble strategies (as shown in this paper). In addition, variability across independent runs may help identify uncertain or ambiguous cases, supporting more transparent validation through comparison of model rationales, confidence estimates, and chromatographic evidence.

Several limitations of the current framework should be considered. First, representing high-resolution chromatographic data as numerical arrays incurs substantial token usage, leading to increased computational cost and latency during inference. Second, model performance depends in part on prompt structure and the clarity with which domain knowledge is encoded, highlighting the need for standardized prompting strategies or systematic prompt optimization. Third, although zero-shot reasoning enables broad applicability, further improvements may be achieved through domain-specific fine-tuning or hybrid approaches that combine learned representations with reasoning-based inference.

Despite these limitations, ChatDIA can directly benefit from ongoing advances in general-purpose LLM capabilities. Improvements in model efficiency, context handling, and reasoning performance are likely to translate into immediate gains in analytical accuracy and scalability without requiring redesign of the underlying workflow. This contrasts with conventional purpose-built tools, which often require retraining or reengineering to incorporate methodological advances.

Beyond DIA peak group selection, the presented approach could be extended to a wide range of mass spectrometry-related tasks, including quality control, interference detection, retention time alignment, and method optimization. More broadly, this work highlights the potential of LLM-based systems to participate directly in structured scientific reasoning over experimental data. Rather than serving solely as tools for pattern recognition or score prediction, such systems can evaluate evidence, compare alternative interpretations, and generate explicit, inspectable rationales linking data to conclusions.

As computational methods continue to evolve, the integration of reasoning-based approaches into scientific workflows may represent a natural progression beyond traditional statistical learning. While current systems remain constrained and require expert validation, their ability to operate on structured data and produce interpretable, evidence-based decisions suggests a future in which computation contributes more directly to the logic of scientific inference. In this context, ChatDIA represents an early example of how such reasoning-driven frameworks can be applied to complex experimental data analysis.

## Conclusion

We conclude that general-purpose LLMs can analyze complex DIA mass spectrometry data and evaluate chromatographic evidence through structured reasoning. ChatDIA highlights the potential of reasoning-based, interpretable AI agents for computational proteomics, particularly in settings where uncertainty, signal ambiguity, and human-in-the-loop interpretation are central.

## Acknowledgement

We thank Dr. John Jumper (DeepMind) for insightful feedback on an early version of this work, particularly regarding the assessment of potential data leakage during model pretraining. This work was supported by the Canadian Institutes of Health Research (CIHR).

---

**Supplementary Figure 1.**
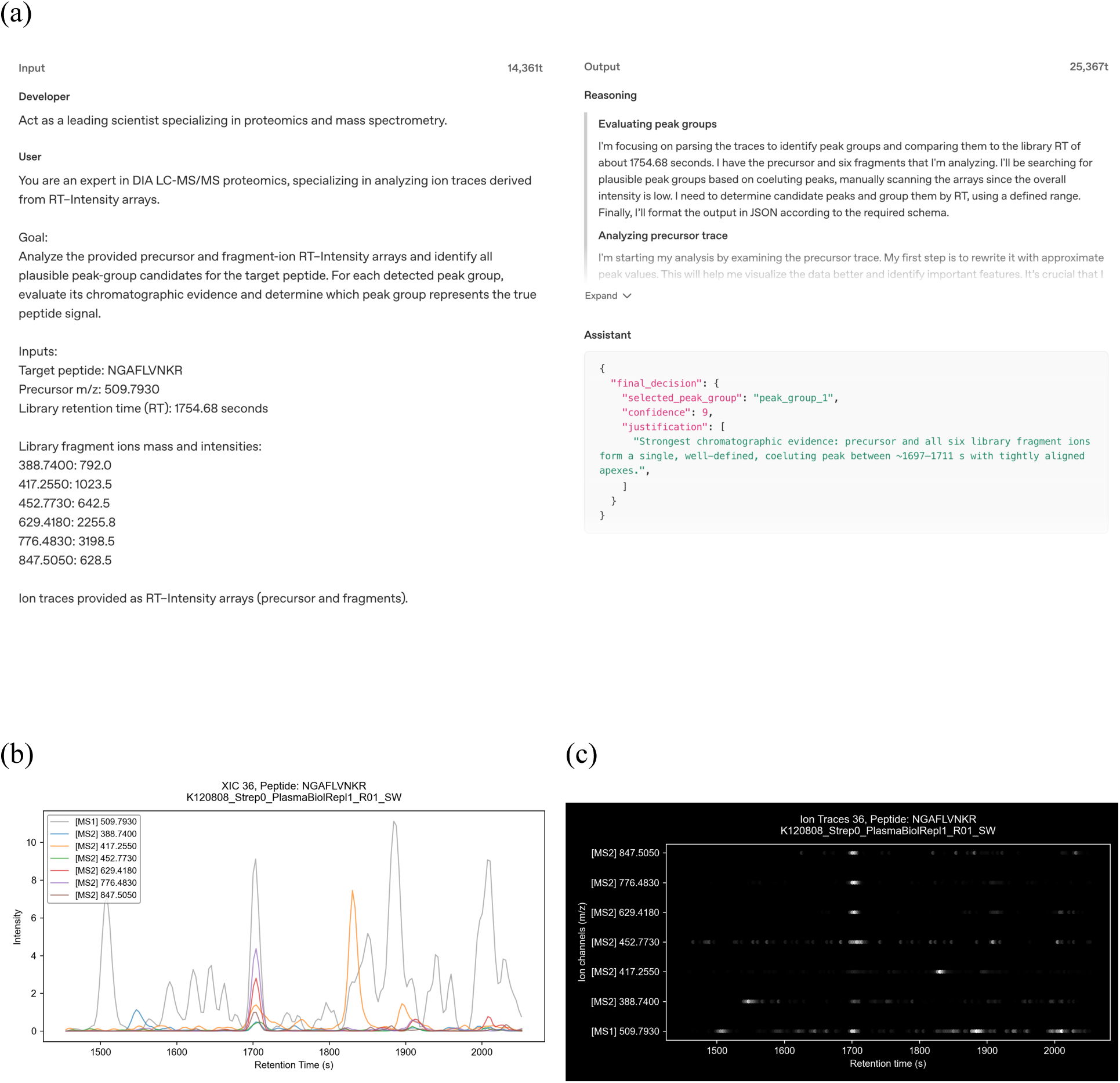
ChatDIA input and output examples. **(a)** Example subset of a ChatDIA prompt and corresponding output generated using GPT-5.2 for a peptide from the *Streptococcus pyogenes* proteomics dataset, including excerpts from the LLM reasoning component and the final decision output. The screenshot is taken from the OpenAI API platform, which stores inputs and outputs from automated ChatDIA analyses. **(b)** Example XICs for a single peptide (transition group), showing precursor and fragment retention time (RT) and intensity information after Gaussian smoothing. Precursor intensities are normalized to fragment intensities to improve visualization. **(c)** Example ion trace image representation for a single peptide (transition group), in which pixel brightness (white intensity) reflects signal strength.

**Supplementary Figure 2.**
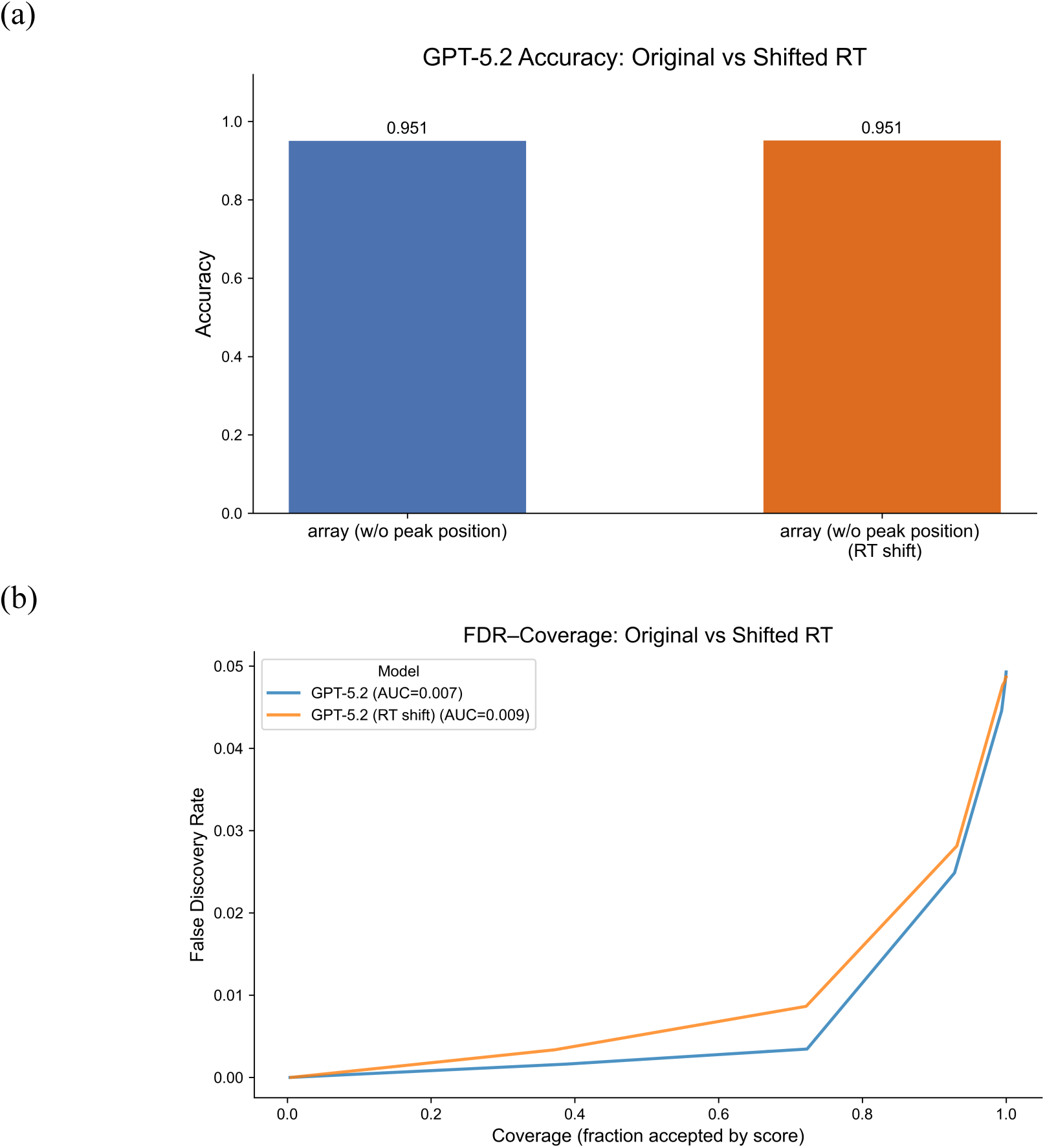
Prior public availability of the *Streptococcus pyogenes* proteomics dataset and potential pretraining exposure do not measurably affect ChatDIA performance. To assess whether prior public availability of the *Streptococcus pyogenes* dataset and potential exposure during GPT pretraining influence model performance, ChatDIA with GPT-5.2 was evaluated under two input conditions: (i) peptide sequence provided with original retention time (RT) arrays, and (ii) peptide sequence masked and with RT arrays shifted by +200 seconds. Performance was evaluated on 423 unique transition groups measured across four technical replicates (1,603 total evaluations). All ChatDIA configurations used numerical XIC array inputs with explicit guidance, but without peak position information. **(a)** Accuracy comparison between the two conditions. **(b)** Risk-coverage (FDR-coverage) analysis comparing performance between the two conditions. Metrics and evaluation procedures are as described in Figure 4.

**Supplementary Figure 3.**
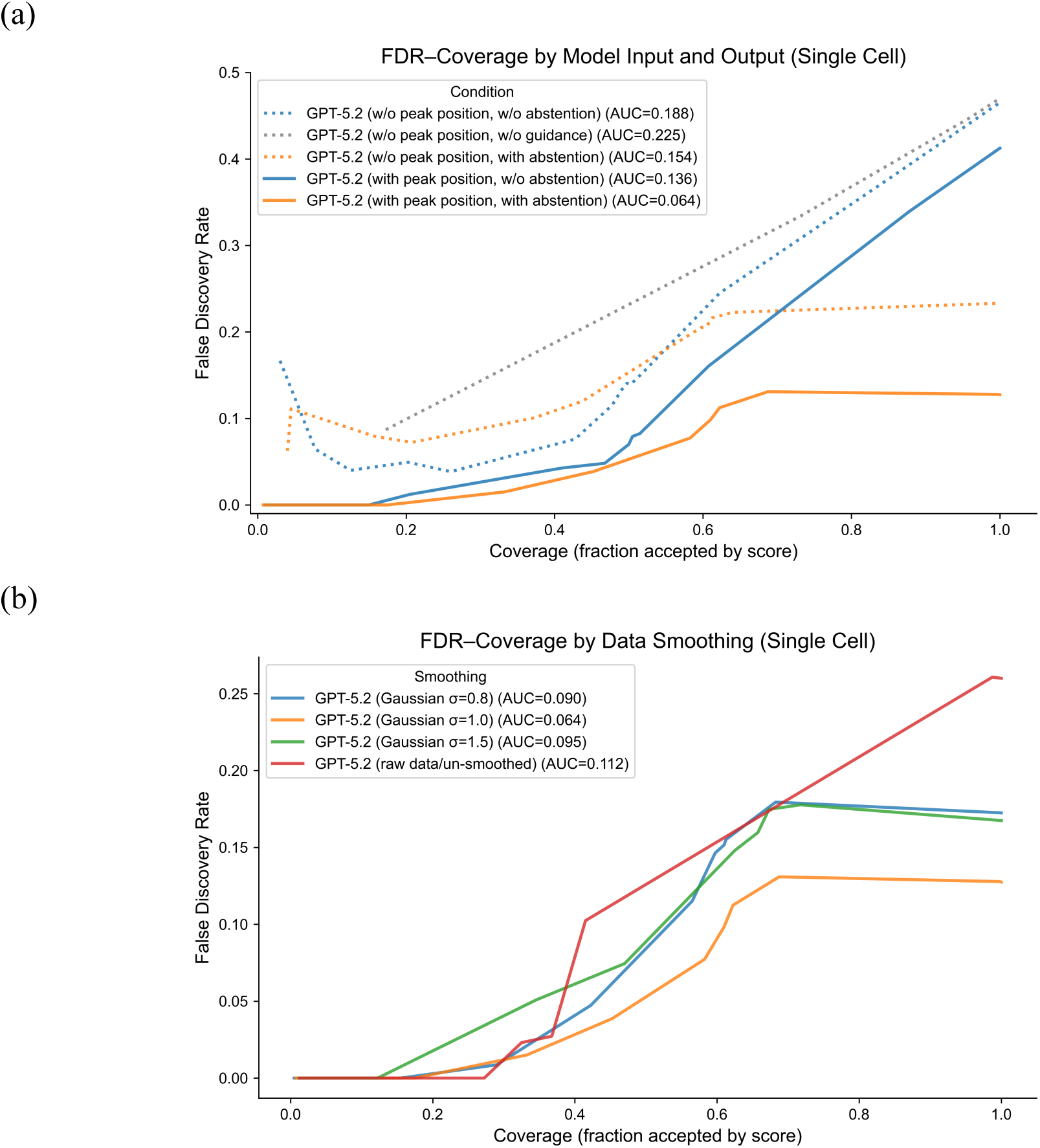
FDR-coverage analysis of ChatDIA under different prompting strategies and data smoothing strengths on a single-cell HEK-293T proteomics dataset. All analyses were performed on the set of 400 manually annotated transition groups. **(a)** FDR-coverage analysis of ChatDIA with GPT-5.2 using numerical XIC array representations with Gaussian smoothing (σ = 1.0). Five conditions are compared: four configurations with explicit guidance, comprising combinations with and without candidate peak group positions provided as input and with and without allowing abstention (i.e., reporting no peak group), and one configuration without candidate peak group information and without explicit guidance. **(b)** FDR-coverage analysis of ChatDIA with GPT-5.2 using numerical XIC array representations incorporating candidate peak group positions as input and allowing abstention. Performance is evaluated across different levels of XIC smoothing, including no smoothing (raw data) and Gaussian smoothing with σ = 0.8, 1.0, and 1.5.

**Supplementary Figure 4.**
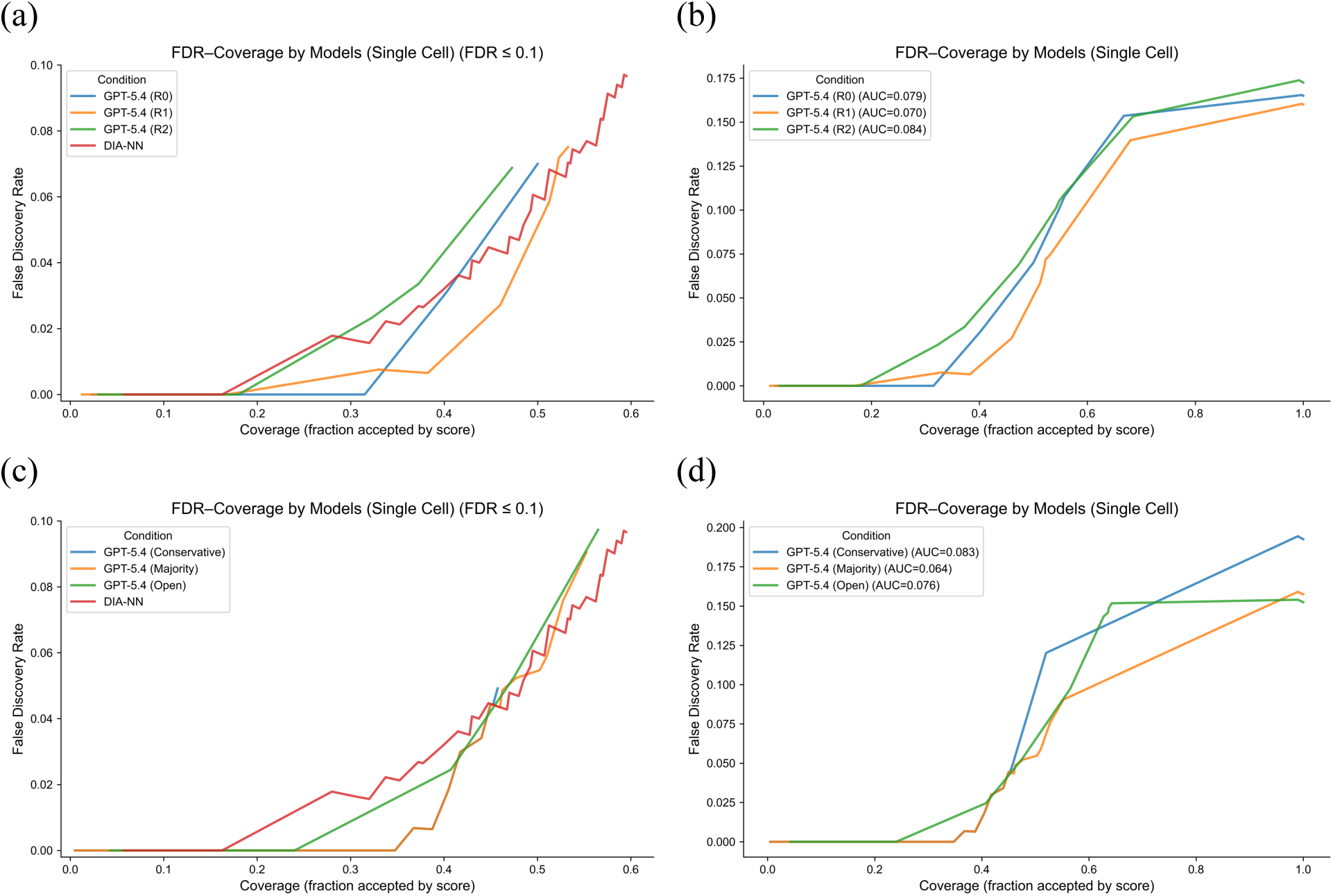
Performance of individual and ensemble LLM runs for DIA proteomics on a single-cell dataset. (a-b) Risk-coverage analysis comparing ChatDIA with GPT-5.4 across three independent LLM runs (R0, R1, and R2), and **(c-d)** risk-coverage analysis comparing ensemble strategies (conservative, majority voting, and open) applied to these runs on a single-cell HEK-293T proteomics dataset. Panels (a) and (c) include DIA-NN as a reference and are shown with zoomed views focusing on the FDR ≤ 0.1 region for improved visualization. Experimental setup, model configuration, and metric definitions are as described in Figure 5.

**Supplementary Figure 5.**
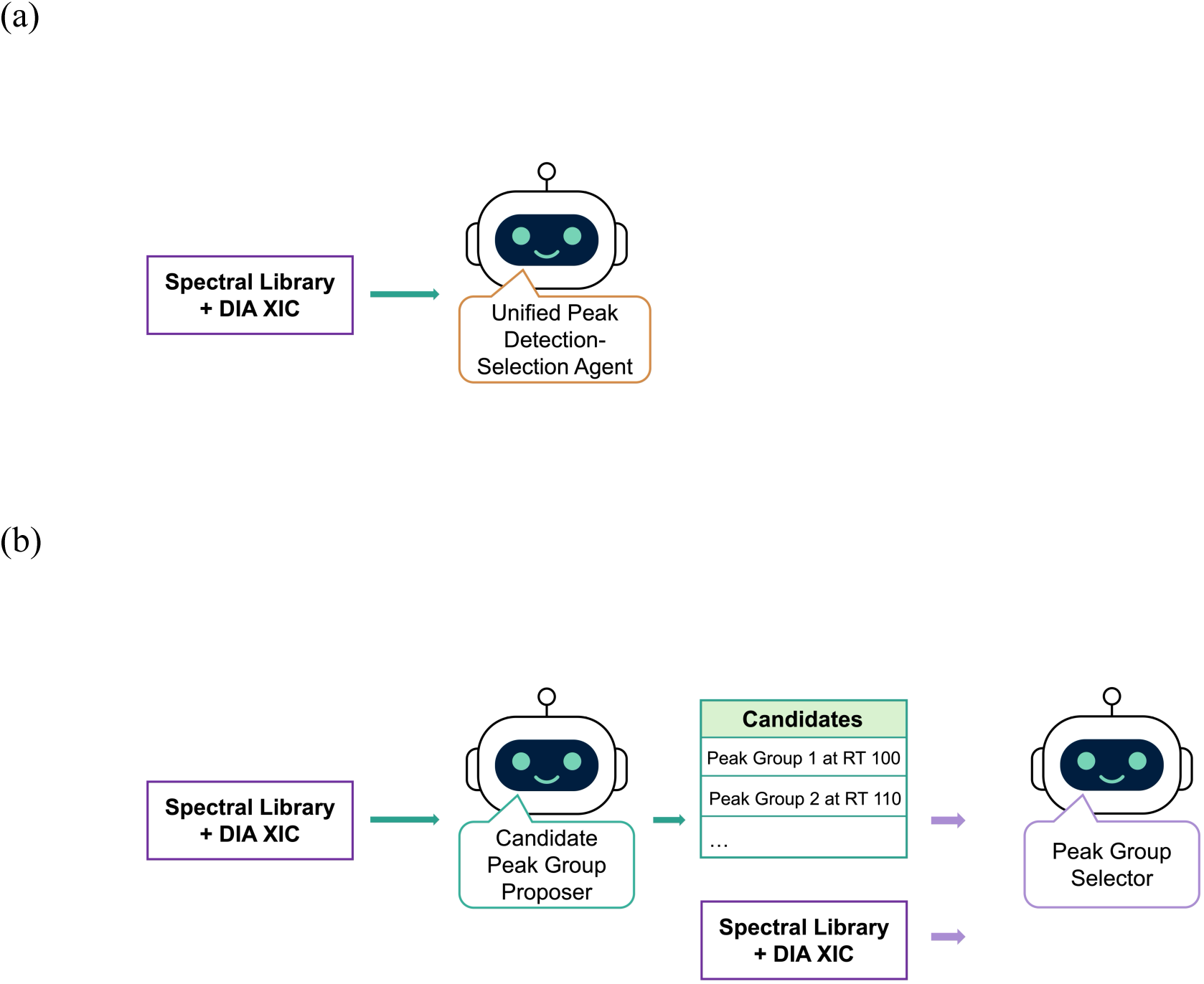
Single-agent and multi-agent ChatDIA workflows for direct peak group detection and selection. **(a)** Single-agent ChatDIA workflow, in which a Unified Peak Detection-Selection Agent performs both candidate peak group detection and selection from DIA XICs, using assay library metadata as reference information. **(b)** Multi-agent ChatDIA workflow, in which a Candidate Peak Group Proposer first identifies candidate peak groups from DIA XICs, followed by a Peak Group Selector that selects the most likely true peptide signal.

**Supplementary Figure 6.**
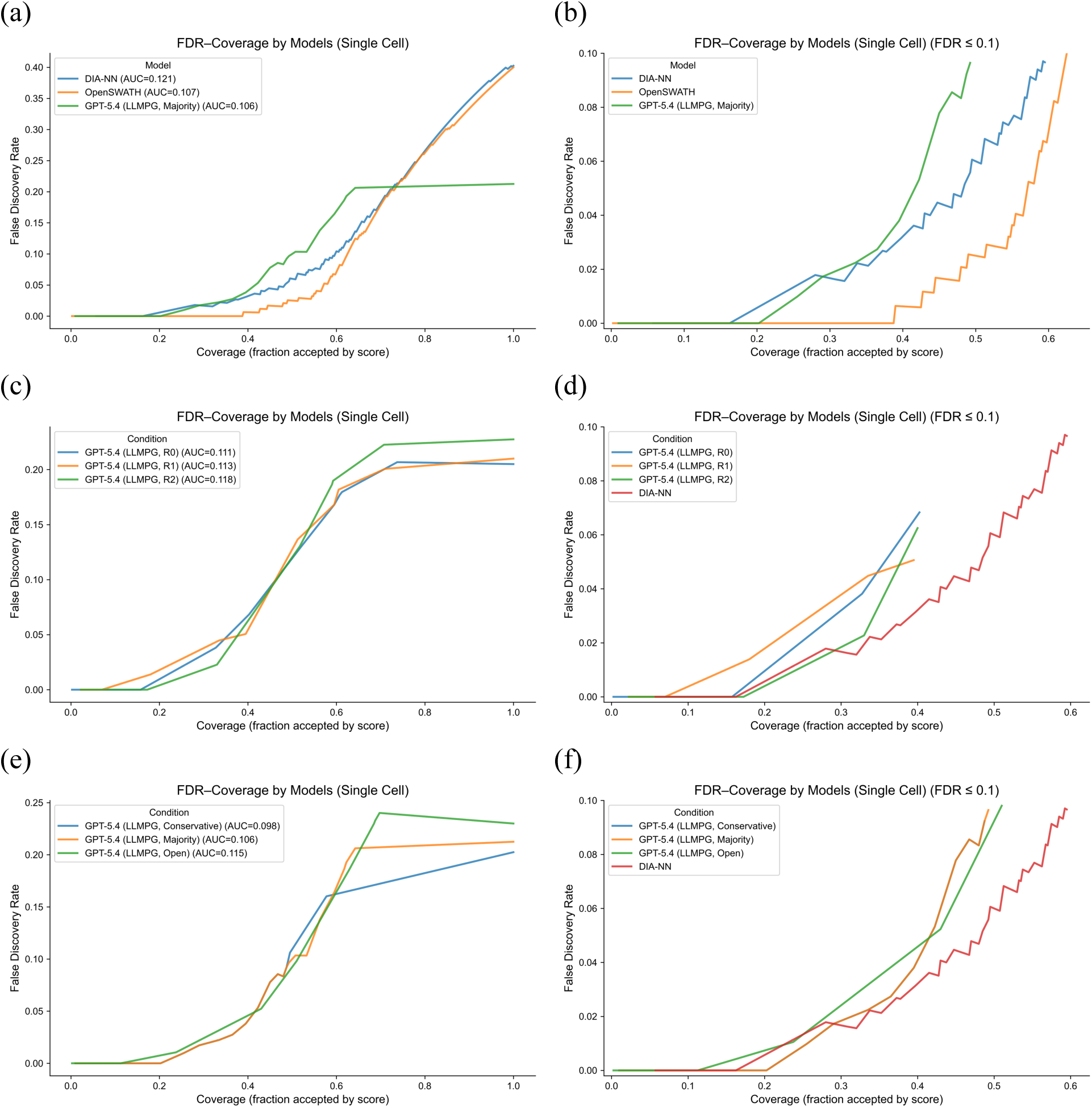
Performance comparison of agentic LLMs and classical algorithms for DIA proteomics on a single-cell dataset. For ChatDIA, a peak-group proposal LLM agent was first used to nominate up to five candidate peak groups, followed by a peak-group selection LLM agent for signal selection and scoring. “LLMPG” in the figure denotes the use of LLM-proposed peak groups. Numerical XIC arrays with Gaussian smoothing (σ = 1.0) were used as input for both agents, and abstention was permitted for the selection agent. **(a-b)** Risk-coverage analysis comparing ChatDIA using GPT-5.4, implemented as an ensemble of three independent runs with majority voting, with OpenSWATH and DIA-NN on a single-cell HEK-293T proteomics dataset. **(c-d)** Risk-coverage analysis comparing three independent ChatDIA runs, denoted R0, R1, and R2. **(e-f)** Risk-coverage analysis comparing ensemble strategies, including conservative, majority-voting, and open strategies, applied to the three runs. DIA-NN is included as a reference in panels **(d)** and **(f)**. Panels **(a)**, **(c)**, and **(e)** show risk-coverage curves over the full operating range, whereas panels **(b)**, **(d)**, and **(f)** show zoomed views focusing on FDR ≤ 0.1. Experimental setup and metric definitions are the same as those used for the Figure 5 experiments.

**Supplementary Figure 7.**
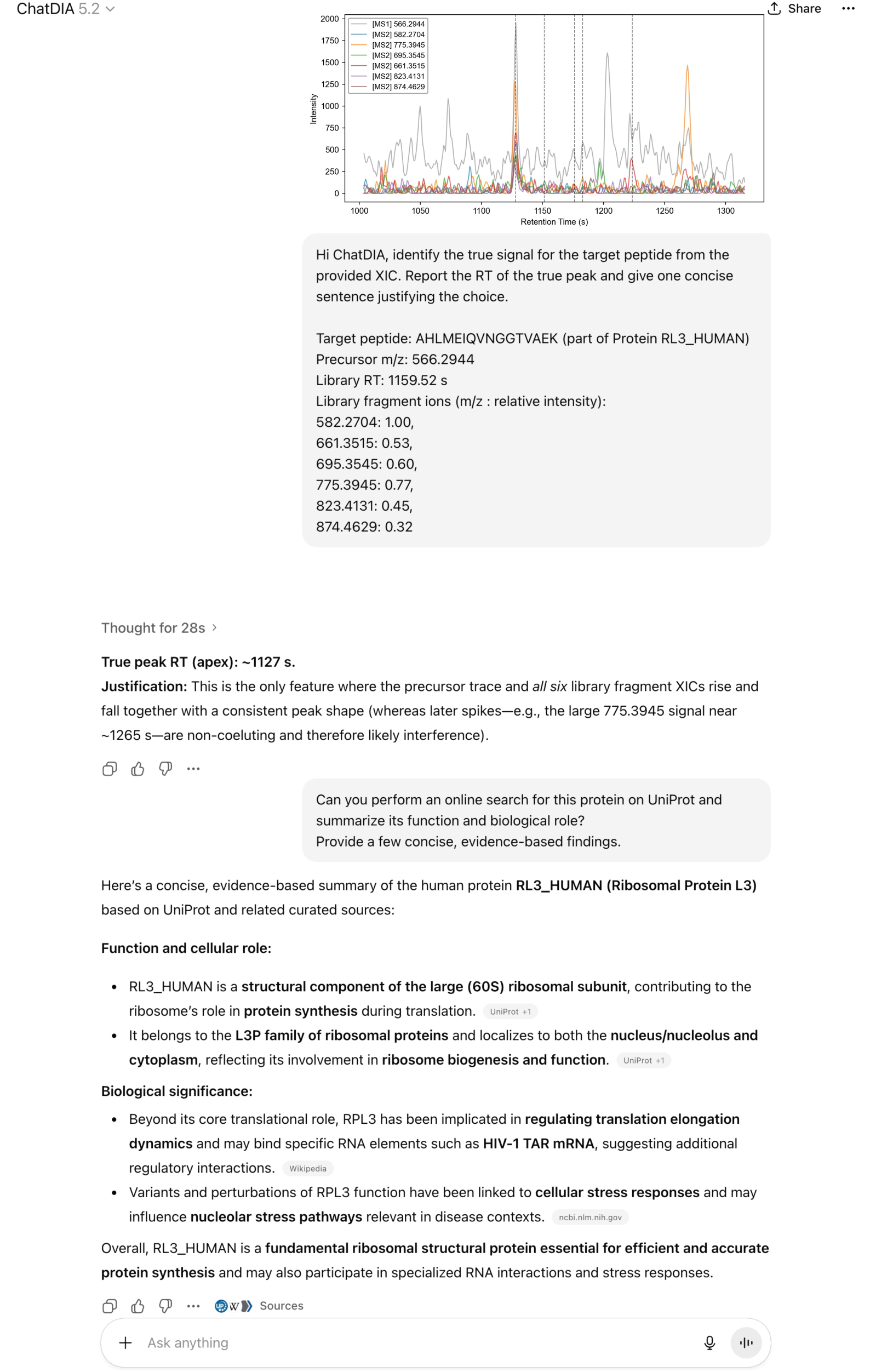
Example use case of ChatDIA. Illustration of ChatDIA usage in a lightweight mode implemented within the OpenAI ChatGPT platform. The example demonstrates how users can interact with the model using natural language to perform DIA analyses, including peptide signal identification by providing an XIC image and assay library metadata. Users can also request ChatDIA to conduct online literature searches to retrieve protein functional information. The peptide shown originates from a single-cell HEK-293T proteomics dataset. The screenshot is taken from the OpenAI ChatGPT platform, where the current lightweight version of ChatDIA is deployed.

**Supplementary Table 1.** Quantitative metrics underlying FDR-coverage analysis for ChatDIA. This table reports performance metrics for ChatDIA with GPT-5.2 using numerical XIC array inputs without peak position information on the *Streptococcus pyogenes* proteomics dataset. Each row corresponds to using the reported LLM confidence score as the decision threshold. For each score-derived threshold, the table reports the resulting coverage, precision, FDR, recall, and the number of model predictions (identified precursor count). Coverage is defined as the fraction of predictions with confidence scores greater than or equal to the threshold, (TP + FP) / (TP + FP + TN + FN), where TP + FP corresponds to the number of predicted precursors above the threshold. Accordingly, the identified precursor count is equivalent to TP + FP and equals coverage multiplied by the total number of evaluated precursors (N = 1,603 transition group evaluations across four technical replicates). Precision is defined as TP / (TP + FP), FDR as FP / (TP + FP), and recall as TP / (TP + FN).

| Score | Coverage | Precision | FDR | Recall | Identified precursor count |
| --- | --- | --- | --- | --- | --- |
| 10 | 0.37% | 1.0000 | 0.0000 | 0.0039 | 6 |
| 9 | 38.68% | 0.9984 | 0.0016 | 0.4062 | 620 |
| 8 | 72.30% | 0.9965 | 0.0035 | 0.7579 | 1159 |
| 7 | 92.83% | 0.9751 | 0.0249 | 0.9521 | 1488 |
| 6 | 99.38% | 0.9554 | 0.0446 | 0.9987 | 1593 |
| 5 | 99.88% | 0.9519 | 0.0481 | 1.0000 | 1601 |
| 4 | 100.00% | 0.9507 | 0.0493 | 1.0000 | 1603 |

**Supplementary Table 2.** Coverage at fixed false discovery rate thresholds for model comparison on a single-cell proteomics dataset. This table summarizes model coverage at fixed FDR thresholds for ChatDIA and established DIA analysis algorithms on a single-cell HEK-293T proteomics dataset. Coverage values are reported at FDR ≤ 0.01, ≤ 0.05, and ≤ 0.10 for DIA-NN, OpenSWATH, and ChatDIA configurations based on GPT-5.4 and GPT-5.2. GPT-5.4 results include individual runs and ensemble strategies across multiple independent LLM runs using Gaussian-smoothed XIC inputs (σ = 1.0), whereas GPT-5.2 results correspond to single-run evaluations with both smoothed (σ = 1.0) and unsmoothed XIC inputs. ChatDIA models use numerical XIC array representations incorporating candidate peak group positions and allow abstention (i.e., reporting no peak group). Coverage is defined as the fraction of predictions with confidence scores greater than or equal to a given score threshold among the 400 manually annotated transition groups (coverage = (TP + FP) / (TP + FP + TN + FN)), where the threshold is determined by method-specific confidence measures (LLM confidence scores for ChatDIA, 1−q values for DIA-NN, and discriminative scores for OpenSWATH). FDR is defined as the fraction of model predictions rejected by manual validation, FP / (TP + FP). For each method and configuration, the reported coverage corresponds to the maximum coverage achievable while satisfying the specified FDR threshold.

| Model | Coverage@FDR≤0.01 | Coverage@FDR≤0.05 | Coverage@FDR≤0.10 |
| --- | --- | --- | --- |
| DIA-NN | 16.25% | 48.00% | 59.50% |
| OpenSWATH | 42.50% | 57.00% | 62.50% |
| GPT-5.4<br>(conservative) | 38.75% | 45.75% | 45.75% |
| GPT-5.4<br>(majority) | 38.75% | 46.25% | 55.25% |
| GPT-5.4<br>(open) | 24.00% | 40.75% | 56.50% |
| GPT-5.4 (R0) | 31.50% | 40.25% | 50.00% |
| GPT-5.4 (R1) | 38.25% | 46.00% | 53.25% |
| GPT-5.4 (R2) | 18.00% | 37.25% | 47.25% |
| GPT-5.2 | 17.50% | 45.25% | 61.00% |
| GPT-5.2<br>(unsmoothed) | 27.25% | 36.75% | 36.75% |

**Supplementary Table 3.** Ensemble strategies for aggregating outputs from multiple independent LLM runs in DIA proteomics. Summary of three ensemble strategies (conservative, majority voting, and open) used to combine peak-detection outputs from three independent runs of the LLM on each transition group in the single-cell HEK-293T dataset. These runs represent repeated LLM inferences on the same input data. Each strategy defines criteria for peak detection based on the number of LLM runs reporting a peak (*support*) and on retention time (RT) agreement, together with corresponding rules for aggregating RT boundaries and confidence scores. In the conservative strategy, a peak is reported only if all LLM runs detect a peak with overlapping RT intervals, yielding a consensus RT defined by interval intersection and a confidence score given by the mean score across runs. In the majority-voting strategy, a peak is reported when at least ⌈N/2⌉ LLM runs detect a peak with overlapping RT intervals, with RT boundaries defined by the median of supporting runs and confidence scaled by the fraction of supporting runs (*support/N*). In the open strategy, the highest-confidence detected peak is selected when at least one LLM run reports a peak. Across all strategies, failure to satisfy the peak-detection or RT-overlap criteria results in no peak assignment and a penalized confidence score defined as the minimum score across runs. Here, *support* denotes the number of LLM runs that report a peak for a given transition group, and *score_support* refers to the confidence scores from those runs. RT overlap is defined as a non-empty intersection of RT intervals, and peak detection requires both RT boundaries to be present.

| Ensemble Strategy | Condition | Ensemble Call | RT Window Rule | Score Rule |
| --- | --- | --- | --- | --- |
| Conservative | All runs detect a peak and RT windows overlap | Peak | RT intersection | $\text{mean}(\text{score}_i)$ |
| | All runs detect a peak but RT windows do not overlap | No peak | -- | $\text{min}(\text{score}_i)$ |
| | At least one run does not detect a peak | No peak | -- | $\text{min}(\text{score}_i)$ |
| Majority voting | $\geq [N/2]$ runs detect peaks and detected runs have RT overlap | Peak | $\text{median}(\text{RT\_left}_i), \text{median}(\text{RT\_right}_i)$ of detected runs | $(\text{support}/N) \times \text{mean}(\text{score\_support})$ |
| | $\geq [N/2]$ runs detect peaks but no RT overlap | No peak | -- | $\text{min}(\text{score}_i)$ |
| | $< [N/2]$ runs detect peaks | No peak | -- | $\text{min}(\text{score}_i)$ |
| Open approach | At least one run detects a peak | Peak | RT from highest-scoring detected run | $\text{max}(\text{score\_support})$ |
| | No runs detect a peak | No peak | -- | $\text{min}(\text{score}_i)$ |

**Supplementary Table 4.** Coverage at fixed false-discovery-rate thresholds for multi-agent and single-agent ChatDIA on a single-cell proteomics dataset. This table summarizes model coverage at fixed FDR thresholds for two ChatDIA configurations: multi-agent ChatDIA, in which one LLM agent was used for candidate peak-group proposal and a second LLM agent was used for peak-group selection, and single-agent ChatDIA, in which a single LLM performed both detection and selection. Both configurations allowed the LLM to abstain from making a peak-group selection. Coverage values are reported at FDR ≤ 0.01, ≤ 0.05, and ≤ 0.10. GPT-5.4 was used, and Gaussian-smoothed XIC arrays (σ = 1.0) were provided as inputs. Results include individual runs and ensemble strategies across multiple independent runs. Coverage is defined as the fraction of predictions with confidence scores greater than or equal to a given score threshold among the 400 manually annotated transition groups (coverage = (TP + FP) / (TP + FP + TN + FN)), where the threshold is determined by method-specific confidence measures (LLM selection confidence scores for ChatDIA). FDR is defined as the fraction of model predictions rejected by manual validation, FP / (TP + FP). For each method and configuration, the reported coverage corresponds to the maximum coverage achievable while satisfying the specified FDR threshold.

| Model | Coverage@FDR≤0.01 | Coverage@FDR≤0.05 | Coverage@FDR≤0.10 |
| --- | --- | --- | --- |
| ChatDIA<br>(multi-agent, R0) | 15.75% | 32.75% | 40.25% |
| ChatDIA<br>(multi-agent, R1) | 7.00% | 33.50% | 39.50% |
| ChatDIA<br>(multi-agent, R2) | 17.25% | 33.00% | 40.00% |
| ChatDIA<br>(multi-agent, conservative) | 25.50% | 39.50% | 48.75% |
| ChatDIA<br>(multi-agent, majority) | 25.50% | 39.50% | 49.25% |
| ChatDIA<br>(multi-agent, open) | 11.25% | 23.75% | 51.00% |
| ChatDIA<br>(single-agent, R0) | 0 | 23.50% | 40.75% |
| ChatDIA<br>(single-agent, R1) | 3.75% | 33.00% | 40.00% |
| ChatDIA<br>(single-agent, R2) | 16.25% | 23.50% | 34.50% |
| ChatDIA<br>(single-agent, conservative) | 14.25% | 35.75% | 37.50% |
| ChatDIA<br>(single-agent, majority) | 3.75% | 32.25% | 46.75% |
| ChatDIA<br>(single-agent, open) | 0 | 31.75% | 44.00% |

